# Achiral Odd Mechanics in Cell Monolayers

**DOI:** 10.64898/2026.07.25.740723

**Authors:** Sreejith Santhosh, Mattia Serra

## Abstract

Cell monolayers are active, orientationally ordered materials whose mechanics can depart from near-equilibrium behavior. Existing continuum theories often inherit assumptions from equilibrium liquid-crystal physics, motivating explicit nonequilibrium formulations. Here, we develop a minimal continuum model of two-dimensional monolayers based on odd mechanics, accounting for broken symmetries and nonequilibrium dynamics. We show that nematic order can support an odd viscous modulus generated by broken time-reversal symmetry and spatial anisotropy, without chirality. The model reproduces half-integer defect motion and stress profiles in Madin–Darby canine kidney (MDCK) monolayers, as well as defect-associated cell accumulation and depletion in neural progenitor and ovarian mesothelium systems. Finally, we estimate the viscous-moduli tensor from measured stress, velocity, and orientation fields in MDCK monolayers and identify a nonzero odd modulus. Our results show that odd mechanics provides a minimal framework for nonequilibrium cell monolayers, complementing conventional active-nematic theories.

## I. INTRODUCTION

Cell monolayers are central to organismal physiology and development [1–3]. Recent work has emphasized how mechanics coordinates cell behavior and how cell behavior, in turn, shapes emergent tissue-scale properties [4–7]. These mechanical features often persist over length and time scales far exceeding those of individual cells [4, 8–10]. They are naturally described by hydrodynamic continuum theories of tissue mechanics and orientational order, which may be polar or nematic [3, 11, 12]. In cell monolayers, polarity and nematicity are typically associated with subcellular biochemical polarity or cell morphology, such as in-plane polarity or cell shape [3]. Here, we focus on monolayers with nematic order. Active nematic models have successfully described diverse cell behaviors, collective migration, and morphogenetic dynamics [13, 14].

Active nematic theory inherits many ingredients from near-equilibrium liquid-crystal physics [15], whose assumptions may not hold in far-from-equilibrium cell monolayers [11]. In particular, continuum formulations often include free-energy-based terms, such as Landau– Ginzburg and orientational-elasticity contributions [14]. Although phenomenologically useful, these terms require care in cell monolayers, which are driven and maintained out of equilibrium [16, 17]. Moreover, predictions of active-nematic theories can depend sensitively on the choice and magnitude of elastic terms [18, 19]. This motivates complementary formulations that retain a continuum description of orientational order while making the nonequilibrium material response explicit.

Here, we formulate a minimal model of monolayer mechanics that incorporates nematic orientational order and explicitly accounts for nonequilibrium material response by building on the recent framework of odd mechanics [20, 21]. This framework provides a continuum approach for describing nonequilibrium mechanical responses across systems ranging from muscle fibers [22] and starfish embryo cell clusters [23] to human crowds [24]. We investigate tissue dynamics generated by a linear viscous constitutive relation with an achiral odd modulus arising from broken spatial and time-reversal symmetries. This framework reproduces experimentally observed motion and stress profiles near half-integer topological defects in tissues. Using velocity and stress data from Madin–Darby Canine Kidney (MDCK) monolayers, we estimate the viscous moduli tensor and identify a nonzero odd modulus. Together, our results show that odd mechanics provides a minimal framework for describing experimentally observed nonequilibrium phenomena in cell monolayers.

## II. RESULTS

### A. Constitutive relations of cell monolayers reveal an achiral odd modulus

We model cell monolayers as a two-dimensional viscous continuum described by a linear constitutive stress strain-rate relation [25–27]. Using the odd-viscosity framework [20, 28], the stress tensor ***σ*** is a linear function of the velocity Jacobian **∇*u***: *σ*_*ij*_ ∝ *C*_*ijkl*_ ∇*u*_*kl*_, where *C*_*ijkl*_ are viscous moduli. The symmetries and conservation laws of the fluid constrain the entries of ***C***. In ordinary fluids, microscopic time-reversal symmetry and Onsager reciprocity imply *C*_*ijkl*_ = *C*_*klij*_ [20]. Cell monolayers, however, are far from equilibrium and can break time-reversal symmetry, allowing antisymmetric components of ***C*** that correspond to nondissipative viscous moduli [28].

Cell monolayers can exhibit uniaxial orientational order, breaking spatial isotropy and allowing anisotropic viscous moduli [13, 29, 30]. Some cell lines also display chirality [31–33]. Here, we focus on achiral cell monolayers, excluding constitutive terms that break in-plane mirror symmetry about the nematic axis. Cytoskeletal contractility, driven by molecular motors, generates an isotropic prestress that contributes to monolayer motion [34–36]. With these modeling assumptions (Fig. 1A), the nondimensional constitutive relation for a cell monolayer takes the form (Sec. S1A for details)

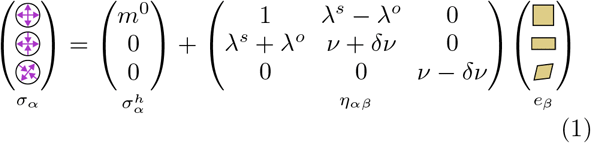

where we decompose ***σ*** and ***e*** = sym(∇***u***) in the matrix basis {***D***^1^, ***D***^2^, ***D***^3^} (Fig. 1B), as in [20, 37]. ***η*** denotes the viscous moduli tensor that maps ***e*** to ***σ*** when both tensors are expressed in the basis ***D***^*i*^. The first basis element, ***D***^1^, captures isotropic deformation, while ***D***^2^ and ***D***^3^ capture the two shear modes of ***σ*** and ***e***. This basis provides a geometric representation of the constitutive relation (Fig. 1B). We omit antisymmetric stress and vorticity components, assuming that cell monolayers are achiral and that stresses are not generated by vorticity. Stress is nondimensionalized by *σ*_*c*_ = *Bv*_*c*_*/x*_*c*_ and strain rate by *v*_*c*_*/x*_*c*_, where *v*_*c*_ is a characteristic velocity, *x*_*c*_ is the length scale of velocity variations, and *B* is the bulk viscosity. The prestress is quantified by *m*^0^ ≥ 0, reflecting monolayer contractility.

**FIG. 1.**
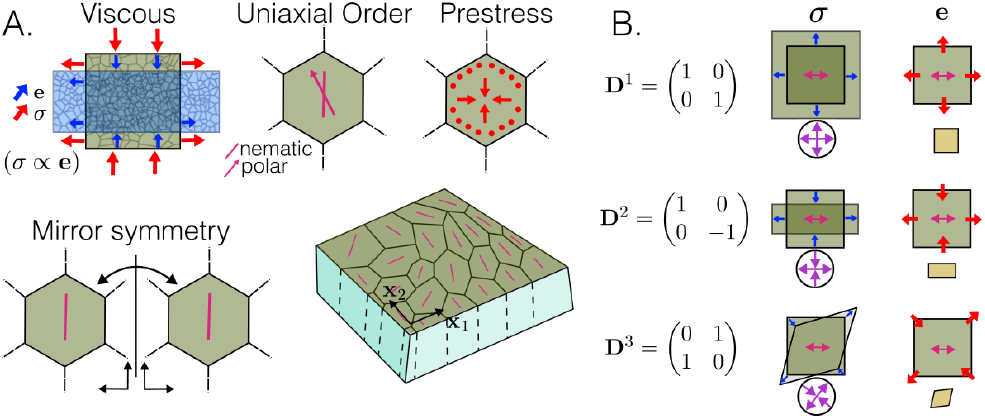
Constitutive relation of cell monolayers: (A) Schematic of the minimal biophysical assumptions. We model cell monolayers as viscous fluids with anisotropy set by uniaxial order. Cell contractility generates an isotropic prestress, while mirror symmetry about the order axis makes the mono-layer achiral. (B) Stress ***σ*** and strain rate ***e*** = sym(**∇*u***) represented in the matrix basis ***D***^*i*^.

Mirror symmetry about the orientational-order axis constrains the allowed components of the viscous-moduli tensor ***η*** (eq. (1)) in the ***D***^*i*^ basis. The odd modulus *λ*^*o*^ arises from broken microscopic time-reversal symmetry and spatial anisotropy induced by orientational order. *λ*^*o*^ nonreciprocally couples the 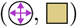, and 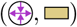 modes (Fig. 1B). The symmetric components of ***η***, i.e. *ν* + *δν, ν* − *δν, λ*^*s*^, are also present in passive anisotropic fluids, while the entries *η*_13_, *η*_31_, *η*_23_, *η*_32_ are forbidden by mirror symmetry about the orientational axis. Although we focus here on the viscous regime, monolayers can also exhibit elastic responses [38, 39]. The odd-mechanics framework can therefore be extended to elastic monolayers, where activity-driven nonconservative microscopic dynamics can generate odd elastic moduli [37]. Most work on odd mechanics has focused on odd moduli arising from broken mirror symmetry, rather than from broken spatial symmetry as derived here. Recent work, however, identified an achiral odd elastic modulus in muscle fibers [22]. Their three-dimensional axisymmetric architecture effectively reduces to a two-dimensional system with orientational order, enabling the nonreciprocal coupling in Eq. (1), but for a stress–strain constitutive relation.

Eq. (1) assumes uniform orientational order, with the nematic axis aligned along **x**_1_ (Fig. 1A). In general, however, orientational order is spatially heterogeneous. We focus on monolayers with nematic order, described by the Q-tensor field ***Q***(**x**, *t*) = 2*S*(***n*** ⊗ ***n*** − I/2), where *S*(**x**, *t*) is the scalar order parameter and ***n***(**x**, *t*) is the director. Thus, Eq. (1) corresponds to the special case *S* = 1 and ***n*** = **x**_1_. For spatiotemporally varying nematic order, and assuming the anisotropic moduli depend linearly on *S*, the viscous-moduli tensor becomes

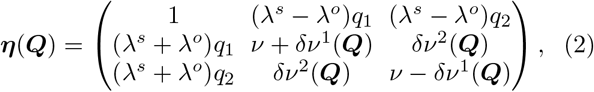

where 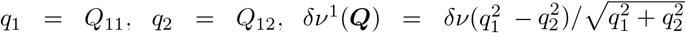 and 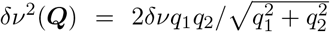 (Sec. S1B for derivation). Because the anisotropic moduli depend linearly on *S*, they vanish for *S* = 0, and Eq. (2) reduces to the constitutive relation of an achiral isotropic fluid. A similar constitutive relation can be derived for monolayers with polar order (Sec. S1B).

### B. The model reproduces observed motion and stress profile around topological defects

To characterize the material response governed by Eq. (2), we compute the steady-state configuration for a prescribed nematic field ***Q***(**x**) by solving **∇** ·***σ*** = 0 with stress-free boundary conditions (Sec. S2). We first analyze topological-defect textures, which are experimentally well characterized and serve as mechanotransduction hotspots in monolayers. A defect of charge *s* at **x** = **0** is defined by ***n***(**x**) = (cos *sθ*, sin *sθ*)^⊤^ and *S*(**x**) = 1 − exp[−*r*^2^/(2*K*^2^)], where *K* is the core size and (*r, θ*) are polar coordinates. We focus on isolated half-integer defects (Fig. 2A), which are common in nematic cell monolayers [13]. Unless stated otherwise, all simulations use a circular domain with *R* = 1 and parameters *K* = 0.05, *m*^0^ = 1, and *ν* = 1. We set *δν* = 0 to isolate the effects of *λ*^*s*^ and *λ*^*o*^; a nonzero *δν* alone does not qualitatively alter the flow fields across nematic textures (Fig. S2A).

**FIG. 2.**
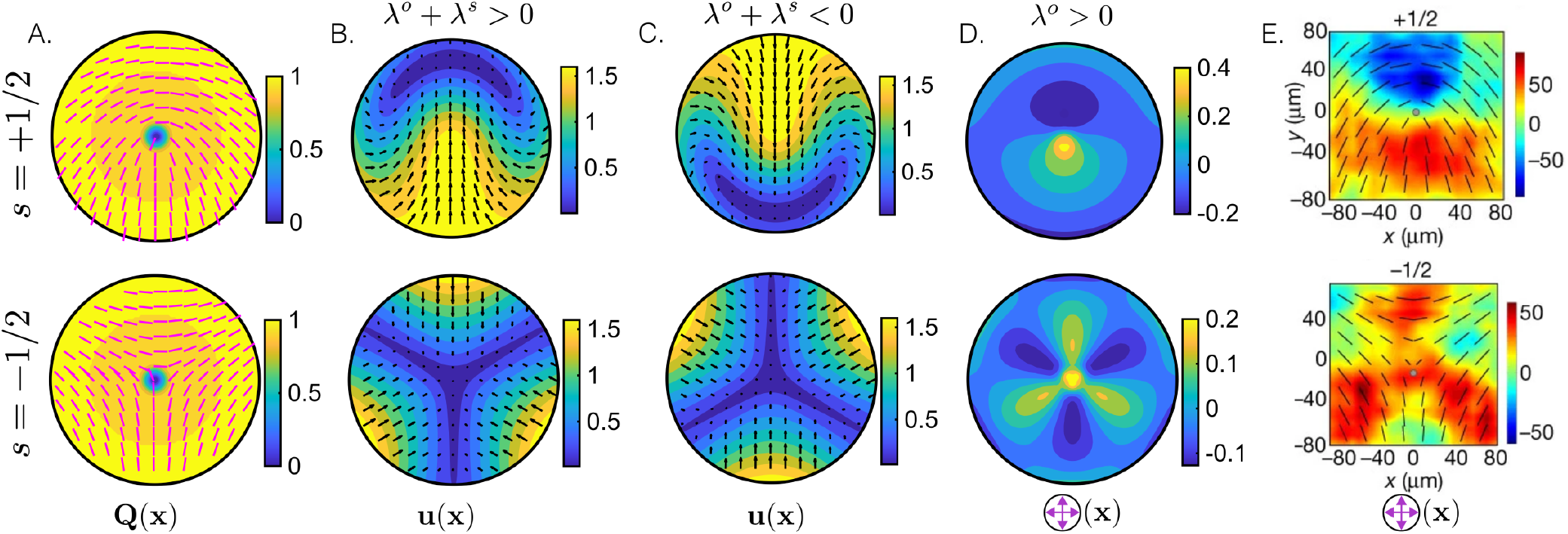
Velocity and stress profile near defects: (A) Nematic tensor field around a topological defect of charge *s*. Color denotes the nematic order parameter *S*(**x**); magenta rods show the director ***n***(**x**). (B–C) Steady-state velocity field **u**(**x**) around defects for *λ*^*o*^ = *λ*^*s*^ = 0.5 (B) and *λ*^*o*^ = *λ*^*s*^ = −0.5 (C), computed from our model. (D) Steady-state isotropic stress 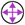(**x**) around defects from our model with *λ*^*o*^ = 1, *λ*^*s*^ = 0. (E) Experimentally inferred 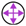(**x**) around half-integer defects in MDCK monolayers, adapted from Fig. 3A in [18]. Stress are in Pa *µ*m. Top and bottom rows correspond to defects with charge *s* = +1/2 and *s* = −1/2.

First, we study the steady-state flow configuration around defects. For *λ*^*o*^ + *λ*^*s*^ > 0, the steady-state velocity configuration around the +1/2 defect exhibits net flow from tail to head at the defect core (Fig. 2B). For *λ*^*o*^ + *λ*^*s*^ < 0, the direction of motion of the +1/2 defect reverses, qualitatively similar to active nematic theories of cell monolayers, where a change from extensile to contractile stress reverses the motion of the +1/2 defect (Fig. 2C). Both tail-to-head and head-to-tail flows have been observed experimentally in cell monolayers, suggesting that either sign of *λ*^*o*^ + *λ*^*s*^ may occur in the same monolayer culture [26, 40]. In contrast, the −1/2 defect does not exhibit a net flow at its core due to its threefold symmetry. For *λ*^*s*^ + *λ*^*o*^ = 0, the flow field around defects becomes radially inwards for both +1/2 and −1/2 defects (Fig. S2A). These results can be understood as the isotropic prestress inducing an isotropic strain rate, which in turn generates an active nematic stress through coupling mediated by the moduli *η*_21_(***Q***) and *η*_31_(***Q***), with strength quantified by *λ*^*s*^ + *λ*^*o*^.

Next, we analyze the isotropic steady-state stress configuration 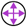(**x**) around defects. Even though the prestress is uniform across the domain, the 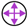(**x**) localizes near the defects (Fig. 2D). The 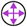(**x**) near both −1/2 and +1/2 defects for *λ*^*o*^ > 0 qualitatively matches the experimentally observed profile in MDCK cell cultures [18] (Fig. 2D-E). The dipolar structure of 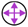(**x**) can be tuned by varying the strength of *λ*^*o*^ (Fig. S2B). This resembles experimentally and theoretically observed changes in stress-localization profiles in active nematic systems upon tuning orientational elasticity [19, 41]. For *λ*^*o*^ > 0, 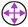(**x**) remains qualitatively consistent with observations in MDCK monolayers. By contrast, regimes with *λ*^*s*^ ≠ 0 and *λ*^*o*^ = 0 fail to reproduce these profiles, particularly around −1/2 defects (Fig. S2C), highlighting the need for nondissipative odd viscosity in cell-monolayer models.

Hydrodynamic active-nematic theories can also explain stress focusing at half-integer defects [18, 19]. However, these theories typically rely on elastic orientational stresses derived from an underlying free energy to generate the correct spatial profiles. The validity of such free-energy dynamics in active nematics remains debated, since cell monolayers are far from equilibrium [17, 42, 43]. Our model avoids this assumption, providing a minimal framework that directly links orientational order to cell-monolayer mechanics.

### C. The model reproduces flow divergence near defects

Next, we examine the steady-state strain-rate profile, specifically 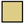(**x**), which affects cell-density dynamics in the monolayer and has implications for crowding and extrusion. We first consider *λ*^*o*^ > 0, the regime consistent with MDCK monolayers, as it reproduces the stress profiles around defects (Sec. II B). In this regime, the model predicts higher compressive strain rate (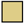 < 0) at the head of a +1/2 defect and at the core of a −1/2 defect relative to the surrounding tissue (Fig. 3A). For *λ*^*o*^ < 0, the 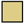(**x**) profiles flip perpendicular to the defect axis for both +1/2 and −1/2 defects (Fig. S2D).

**FIG. 3.**
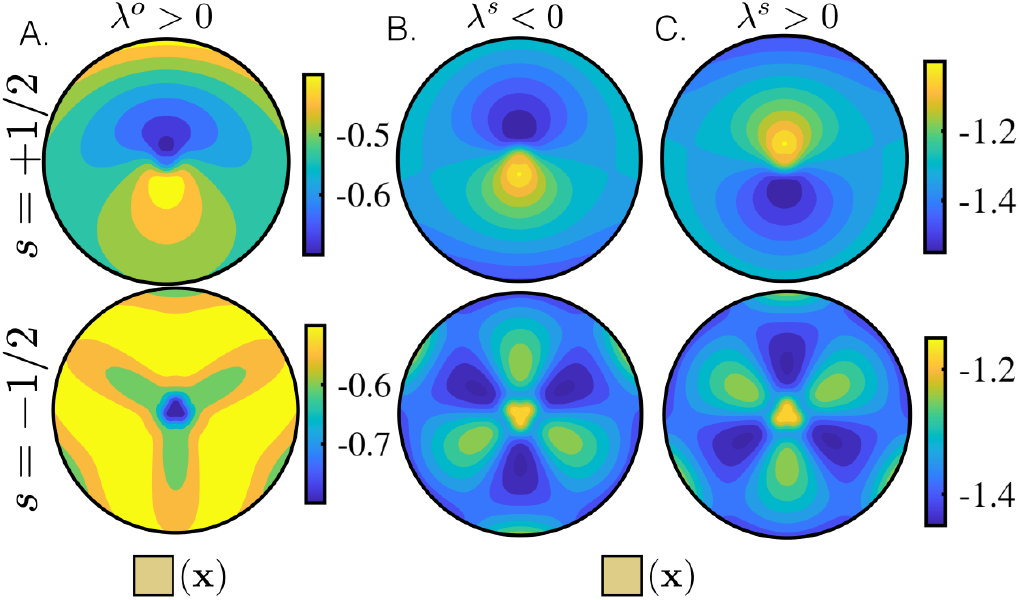
Strain rate profile near defects: (A-C) Steady-state 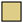(**x**) configuration around a defect using our model, as in Fig. 2. Viscous moduli used are (A) *λ*^*o*^ = 1, *λ*^*s*^ = 0, (B) *λ*^*o*^ = 0, *λ*^*s*^ = −0.5 and (C) *λ*^*o*^ = 0, *λ*^*s*^ = 0.5.

For MDCK monolayers, where *λ*^*o*^ > 0, Ref. [18] showed that extrusion sites spatially correlate with both +1/2 and −1/2 defects. The enhanced extrusion rate near +1/2 defects was attributed to local compressive isotropic stress, 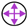 < 0. This mechanism, however, does not apply to −1/2 defects, where the isotropic stress is tensile, 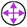 > 0 (cf. Fig. 2D and Fig. 3A in [18]). Our model suggests that compressive strain rate, 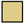 < 0, may also contribute to extrusion near −1/2 defects, where compressive stress is absent at the defect core (Fig. 2D,E). This is consistent with growing evidence that compressive tissue deformation can mediate cell extrusion [44–47]. Thus, our model complements standard hydrodynamic active-nematic theories, which commonly assume incompressibility, and reveals flow-divergence patterns with direct physiological relevance.

Next, we isolate the mechanical effect of *λ*^*s*^ by setting *λ*^*s*^ ≠ 0 and *λ*^*o*^ = 0. For −1/2 defects, this regime produces a local maximum of 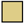(**x**) at the defect core (Fig. 3B), in contrast to the local minimum obtained for *λ*^*o*^ ≠ 0 (Fig. 3A). Around +1/2 defects, the 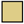(**x**) profile remains dipolar, similar to the *λ*^*o*^ ≠ 0 case (Fig. 3B). Through the continuity equation, 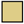(**x**) shapes cell-density dynamics: compressive regions, 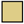 < 0, increase local density, whereas expanding regions, 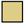 > 0, decrease it. Our model predicts larger accumulation at the head of +1/2 defects for *λ*^*s*^ < 0 and at the tail for *λ*^*s*^ > 0, together with depletion at the −1/2 defect core relative to the surrounding tissue. Similar accumulation–depletion dynamics have been observed in murine neural progenitor cells (NPCs), ovarian mesothelium monolayers, and bacterial colonies by tracking local density changes near defects [48–50]. These observations were previously interpreted through anisotropic substrate friction and active nematic stresses. Here, we show that a linear viscous constitutive relation with anisotropic modulus *λ*^*s*^ can reproduce these density dynamics.

### D. Experimental estimation of *η*

Achirality constrains the entries of ***η*** (Eq. (1)), while the nonequilibrium nature of cell monolayers permits a nonzero odd modulus *λ*^*o*^. To test this assumption and infer *λ*^*o*^, we estimate ***η*** from experimental data. Bayesian Inference Stress Microscopy (BISM) reconstructs the internal stress field ***σ***(**x**, *t*) from traction-force measurements in cell monolayers [51, 52]. Particle Image Velocimetry (PIV) provides the velocity field **u**(**x**, *t*) [53], from which we compute the strain-rate tensor ***e***(**x**, *t*). Assuming a constant prestress, we fit the linear relation between ***σ*** and ***e*** to estimate ***η*** using Eq. (1). Because this constitutive relation is defined in a coordinate system locally aligned with the nematic director ***n***(**x**, *t*), we introduce the local frames 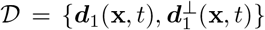, with ***d***_1_(**x**, *t*) = ***n***(**x**, *t*). We take ***n***(**x**, *t*) to align with the cell long axis, obtained from cell-shape segmentation.

We perform this analysis on *N* = 5 independent MDCK datasets from [46], containing spatiotemporal stress, velocity, and cell-segmentation data for monolayers cultured on a 15 kPa substrate. We express both the stress ***σ*** and strain rate ***e*** in the local frame *D*, decompose them in the basis ***D***^*i*^, and infer ***η*** (SI Sec. S3). Because the data are aggregated over space and time, the inferred moduli represent spatiotemporal averages. As described in Sec. II A, we normalize ***η*** by the *η*_11_ entry, corresponding to the bulk viscosity. The entries (*η*_31_, *η*_13_, *η*_32_, *η*_23_) are close to zero, consistent with the assumed achiral symmetry (Fig. 4A). The diagonal shear viscosities *η*_22_ and *η*_33_ are nonzero and not significantly different (Fig. 4B). By contrast, *η*_21_ is nonzero (Fig. 4A) and differs significantly from *η*_12_ (Fig. 4C), implying a nonzero odd-viscous modulus *λ*^*o*^ ≈ 0.1 and reflecting broken Onsager reciprocity in cell monolayers (Fig. 4C, SI Sec. S3). Its positive sign is consistent with our model prediction obtained by matching defect dynamics and stress profiles in MDCK monolayers (Fig. 2).

**FIG. 4.**
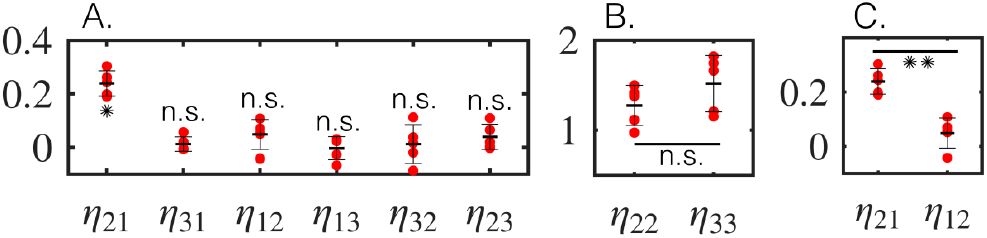
Estimating viscous moduli of MDCK cell monolayers: (A) Non-diagonal entries of the viscous modulus tensor ***η***. ∗ denotes *p* < 10^−3^ and *n*.*s* denotes *p* > 0.05 for a one-sample t-test with target mean of zero. (B) Diagonal entries of ***η***. *n*.*s* denotes *p* > 0.05 for a paired sample t-test. (C) *η*_21_ and *η*_12_. ∗∗ denotes *p* = 10^−2^ for a paired sample t-test. We used *N* = 5 independent measurements of stress and strain rate in MDCK monolayers from [46] (SI Sec. S3 for estimation details).

## III. DISCUSSION

We developed a minimal continuum model of cell-monolayers using the framework of odd mechanics. Odd mechanics provides a continuum description of materials with broken symmetries and nonequilibrium constitutive responses, relaxing the constraints of near-equilibrium constitutive theories. We show that cell monolayers can exhibit an achiral odd modulus arising from spatial anisotropy and nonequilibrium activity. The model further accounts for spatiotemporal variations in orientational order and reproduces observed mechanical response around topological defects. In particular, it captures the stress patterns measured near defects in MDCK monolayers and explains defect-associated density variations across distinct biological systems. The model presented here assumes viscous mechanics with nematic order, but can be readily extended to incorporate polar order and elastic mechanical responses (Sec. S1A, S1B).

Within this framework, mirror symmetry about the local orientational-order axis constrains the viscous-moduli tensor while permitting odd mechanical couplings. We verify these assumptions by inferring the tensor from measured stress, velocity, and orientation fields in MDCK monolayers and find a positive achiral odd modulus, with which our model consistently reproduces the mechanical responses observed near defects.

Recent work has emphasized the role of spatiotemporal heterogeneity in cell-monolayer mechanics [26, 40, 54]. In particular, spatial variations in viscosity have been linked to heterogeneous energy injection [26]. More broadly, odd mechanics offers a route to infer broken symmetries from the structure of the viscous-moduli tensor ***η***. Mapping the spatiotemporal variations of ***η*** may therefore reveal regions of enhanced orientational order, including nematicity and polarity, as well as signatures of activity associated with broken time-reversal symmetry [20].

Another relevant future direction is identifying the microscopic mechanisms underlying these odd moduli [20]. Although likely system-specific, such mechanisms should coarse-grain to the continuum description proposed here. For cell monolayers, cell-resolved vertex and phase-field models provide natural frameworks for determining how cellular processes generate odd mechanical couplings and how these can emerge without explicit chirality or broken mirror symmetry. More broadly, because achiral odd moduli are permitted in active nematic systems, similar responses may arise in cytoskeletal networks, bacterial biofilms, and other active materials with nematic order.

## Acknowledgments

We thank Austin Hopkins for insightful discussions. M.S. acknowledges support from NSF PHY-2413073 and NSF CAREER PHY-2443851, NIH R35GM156889 and HFSP RGEC31/2024.

## Supplementary information

### S1. DERIVATION OF CONSTITUTIVE RELATION FOR CELL MONOLAYERS

#### A. Uniform orientational order

Modeling the mechanical response of a cell monolayer requires a constitutive relation linking stress to material deformation. Although cell monolayers may exhibit both elastic and viscous behavior, here we focus on the viscous regime. We assume a linear relation between the stress ***σ***(***x***, *t*) and the velocity gradients **∇*u***(***x***, *t*), where ***u***(***x***, *t*) = (*u*_1_(***x***, *t*), *u*_2_(***x***, *t*))^⊤^ denotes the velocity field. Because cell monolayers are intrinsically out of equilibrium, we adopt the framework of odd mechanics, which describes materials with broken symmetries and conservation laws [1]. We express the resulting constitutive relation by representing the stress and velocity-gradient tensors in the symmetric matrix basis

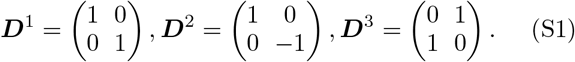

In doing so, we restrict the constitutive description to the symmetric components of these tensors. The stress is taken to be symmetric because we assume the absence of internal torque densities and couple stresses, while the antisymmetric part of **∇*u***, corresponding to local rigid-body rotation or flow vorticity, is assumed not to generate stress. The constitutive relation for a viscous material in the matrix basis {***D***^*i*^} is given by

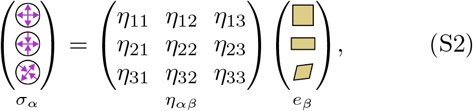

where ***η*** is the viscous modulus tensor allowing for all coupling between ***σ*** and ***e*** = sym(**∇*u***). The viscous-modulus tensor ***η*** is then constrained by the symmetries and conservation laws of the material. For example, the constitutive relation of an isotropic material must remain invariant under coordinate rotations of the form ***x*** → ***R***(*θ*)***x***, where ***R***(*θ*) is the two-dimensional rotation matrix. Under such a transformation, the vectors ***σ*** and ***e*** transform as ***σ*** → ***A***(*θ*)***σ*** and ***e*** → ***A***(*θ*)***e***, where ***A***(*θ*) is given by

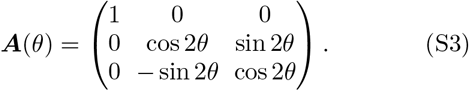

Invariance of the constitutive relation under frame rotations requires ***η*** to transform as ***η*** → ***A***(*θ*)^⊤^***ηA***(*θ*), and therefore imposing the conditions

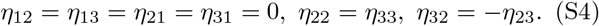

Cell monolayers typically exhibit uniaxial orientational order, which breaks material isotropy. Consequently, the entries of ***η*** are not constrained by Eq. (S4), except in regions where orientational order vanishes (see Sec. S1 B). Here, we assume that the monolayer is achiral, so that the constitutive relation is invariant under reflection about the orientational axis. Taking the uniaxial order orientation aligned with ***x***_1_ (Fig. S1A), this mirror symmetry corresponds to invariance of Eq. (S2) under the transformation {***x***_1_, ***x***_2_}→ {**x**_1_, − **x**_2_}. Under this transformation, the vectors ***σ*** and ***e*** transform as 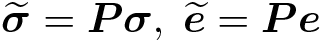, where ***P*** is the parity-transformation

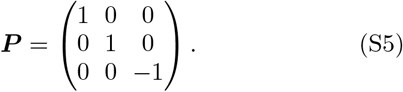

Invariance of Eq. (S2) under parity transformation given in Eq. (S5) enforces the constraint

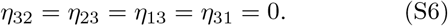

Therefore, for an achiral viscous cell monolayer with uniform orientational order along ***x***_1_, the constitutive relation is constrained to the form

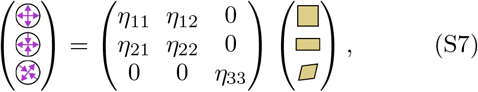

where *η*_11_ is the bulk viscosity and (*η*_22_, *η*_33_) are the shear moduli for two shear modes. The moduli (*η*_21_, *η*_12_) couple the isotropic 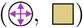 and on-axis shear 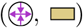 modes. For ordinary fluids, whose microscopic dynamics obey time-reversal symmetry, the viscous-modulus tensor is symmetric, ***η*** = ***η***^⊤^. By the Onsager reciprocal relations, this implies *η*_12_ = *η*_21_. In contrast, the nonreciprocal coupling *η*_12_ ≠ *η*_21_ is permitted only in materials with broken microscopic time-reversal symmetry. Cell monolayers satisfy this condition because they are active, energy-consuming systems operating far from thermodynamic equilibrium.

Cell monolayers are typically prestressed by cell contractility. We incorporate this prestress by modifying the constitutive relation as

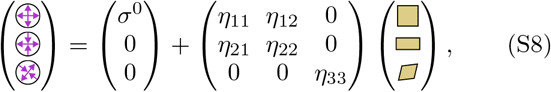

**Figure S1.**
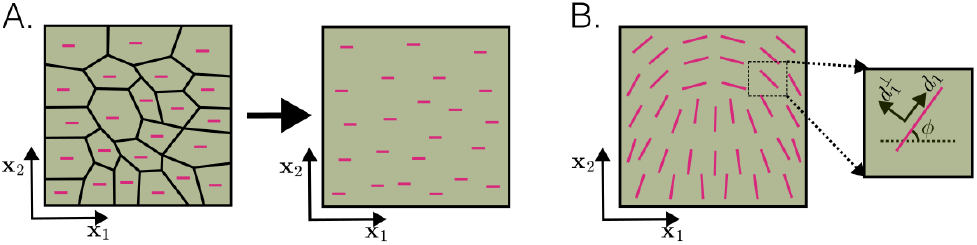
Schematic of orientational order: (A) Cell monolayers are modeled as viscous continua with orientational order, with motion described in the global coordinate frame *G*: {***x***_1_, ***x***_2}_. (B) Schematic of a cell monolayer with spatially varying nematic order. The local frame *D*: {***d***_1_, ***d***_2}_ is aligned with the nematic director at angle *ϕ*(***x***, *t*). The pink arrow indicates the local nematic axis.

where *σ*^0^ quantifies the cell contractility. We next non-dimensionalize Eq. (S8) by rescaling ***e*** with *v*_*c*_*/x*_*c*_ and ***σ*** with *η*_11_*v*_*c*_*/x*_*c*_ where (*v*_*c*_, *x*_*c*_) are the characteristic velocity and length-scale. The non-dimensionalized constitutive relation is given by

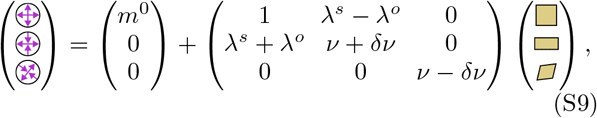

where *m*^0^ = *σ*^0^*x*_*c*_*/η*_11_*v*_*c*_, *λ*^*s*^ = (*η*_21_ + *η*_12_)/(2*η*_11_), *λ*^*o*^ = (*η*_21_ − *η*_12_)/(2*η*_11_), *ν* = (*η*_22_ +*η*_33_)/(2*η*_11_) and *δν* = (*η*_22_ − *η*_33_)/(2*η*_11_). The constitutive equation for an achiral, prestressed and elastic cell monolayer with uniaxial order would follow the same derivation as given above although with the strain-rate replaced with strain and the viscous moduli replaced with the elastic moduli.

#### B. Spatiotemporal orientational order

The constitutive relation derived in Eq. (S9) assumes uniform orientational order. In cell monolayers, however, the orientational order varies in space and time (Fig. S1B). Here, we restrict our analysis to nematic uniaxial order, which has been observed in a variety of cell monolayers. Nematic order is quantified by the Q-tensor field ***Q***(***x***, *t*) = 2*S*(***x***, *t*)(***n***(***x***, *t*) ⊗ ***n***(***x***, *t*) − I/2), where ***n***(***x***, *t*) = [cos *ϕ*(***x***, *t*), sin *ϕ*(***x***, *t*)] is the nematic director axis and *S*(***x***, *t*) is the nematic order parameter. The value *S* = 0 indicates the absence of nematic order, in which case the material is isotropic. Therefore, when *S* = 0, ***η*** must satisfy the constraints in Eq. (S4). Assuming that Eq. (S9) describes the constitutive relation for *S* = 1 and ***n*** = ***x***_1_, we minimally incorporate this constraint by modifying ***η*** in Eq. (S9) as

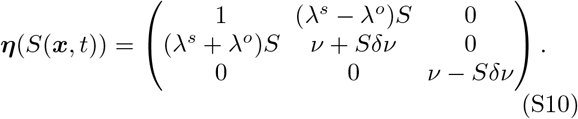

For cell monolayers with spatiotemporal variations in ***n*** but uniform *S*, Eq. (S9) holds locally in the coordinate system 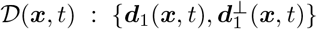, where ***d***_1_(***x***, *t*) is aligned with the local nematic director (Fig. S1B). The constitutive relation in *D*(***x***, *t*) can be expressed in the global frame *G* : {***x***_1_, ***x***_2_} by transforming ***η*** as

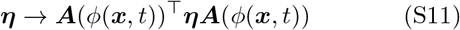

under the rotational coordinate transformation ***A*** (Eq. (S3)) through the angle *ϕ*(***x***, *t*). Accounting for both the spatially varying order parameter in Eq. (S10) and the varying nematic director field in Eq. (S11), the constitutive relation in the global frame *G* is given by

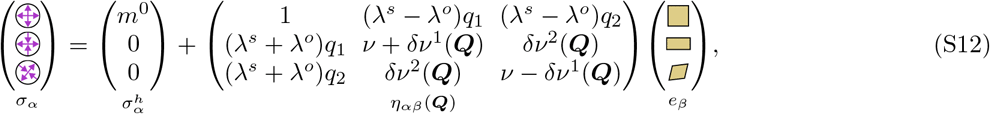

where (*q*_1_, *q*_2_) = (***Q***_11_, ***Q***_12_) and 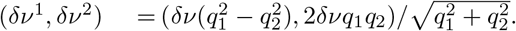.

Cell monolayers also exhibit polar orientational order, which can be quantified using the polar order field ***p***(***x***, *t*) = *p*(cos *ϕ*, sin *ϕ*)^⊤^ [3]. The polar order parameter *p* captures the polar anisotropy of the material, with *p* = 0 corresponding to an isotropic material. Similar to epithelia with nematic order, to model the anisotropy dependence, we assume a linear dependence on the order parameter *p* (*λ*^*s*^, *λ*^*o*^, *δν*) → (*λ*^*s*^*p, λ*^*o*^*p, δνp*). Using these modeling assumptions, the constitutive relation for a cell monolayer with polar order is given by

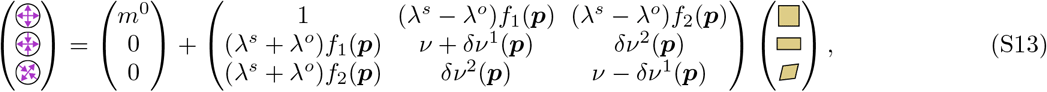

where 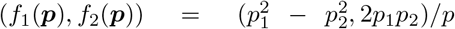 and (*δν*^1^(***p***), *δν*^2^(***p***)) = *δν*(*f*_1_(***p***)^2^ − *f*_2_(***p***)^2^, 2*f*_1_(***p***)*f*_2_(***p***))*/p*.

### S2. MECHANICAL RESPONSE NEAR TOPOLOGICAL DEFECTS

To study the behavior of a material with nonuniform nematic order, we solve for its steady-state configuration for a prescribed nematic field ***Q***(***x***). This configuration is obtained from the overdamped momentum balance

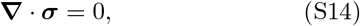

where 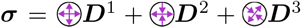. We are particularly interested in the mechanical response near topological defects. The Q-tensor field of a defect with topological charge *s* located at ***x*** = 0 is defined by *ϕ*(***x***) = *sθ* and 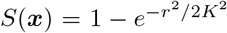 where (*r, θ*) are polar coordinates and *K* denotes the defect core size. Throughout this work, we consider a circular domain of radius *R* = 1 and impose a stress-free boundary condition

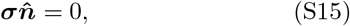

where 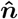 is the normal vector at the boundary. The solution of Eqs. (S14) and (S15) is invariant under rigid-body motions of the domain, and therefore the velocity field is not uniquely determined. To obtain a unique numerical solution, we replace the traction-free boundary condition with the Robin boundary condition

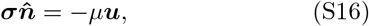

where *µ* is the viscous drag coefficient associated with motion of the monolayer at the boundary. We choose *µ* to be sufficiently small that the boundary remains approximately traction-free. The resulting system of equations in nondimensional form is

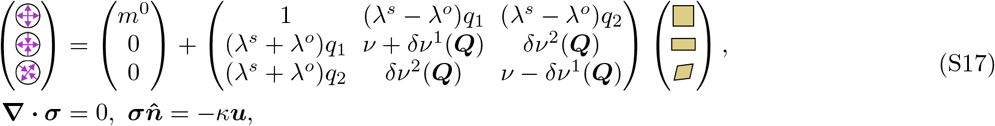

where *κ* = *µx*_*c*_*/η*_11_. We solve Eq. (S17) using the Partial Differential Equations Toolbox in MATLAB [4]. For all simulation results, we verify numerical robustness by varying the spatial discretization. We set *κ* = 10^−4^ and vary it from 10^−5^ to 10^−3^ to establish robustness with respect to their values. We choose *κ* ≪ 1 so that the boundary condition approximates stress-free conditions. Solving Eq. (S17) yields the steady-state velocity field ***u***, from which we compute ***e***. The stress field ***σ*** is then obtained from ***e*** using Eq. (S12).

### S3. ESTIMATING *η* FROM EXPERIMENTS

We analyzed the dataset reported in [2], which contains stress ***σ***, velocity ***u***, and cell long-axis orientation fields from *N* = 5 independent spatiotemporal measurements of MDCK monolayers. The stress field was inferred using Bayesian Inference Stress Microscopy, the velocity field was obtained using Particle Image Velocimetry, and the cell long-axis orientations were extracted from cell segmentation. Assuming that the cell long axis defines the local nematic orientation, we computed the orientational angle field *ϕ*(***x***, *t*). Using *ϕ*(***x***, *t*), we defined the local frame 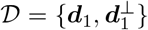, where ***d***_1_ = [cos *ϕ*(***x***, *t*), sin *ϕ*(***x***, *t*)]. In this frame, the constitutive relation is given by Eq. (S8). To estimate ***η***, we computed **∇*u*** from the velocity field, expressed both **∇*u*** and ***σ*** in the local frame *D*, and decomposed them onto the tensor basis ***D***^*i*^. Assuming a spatially uniform *σ*^0^, we fitted the linear relation between ***σ*** and **∇*u*** using MATLAB’s `fitlm` function to estimate ***η***. We then normalized the moduli by *η*_11_ and compared the resulting values with Eq. (S9) to extract *λ*^*o*^ and *λ*^*s*^. One-sample and paired sample *t*-tests were performed using MATLAB’s `ttest` function.

---

**Figure S2.**
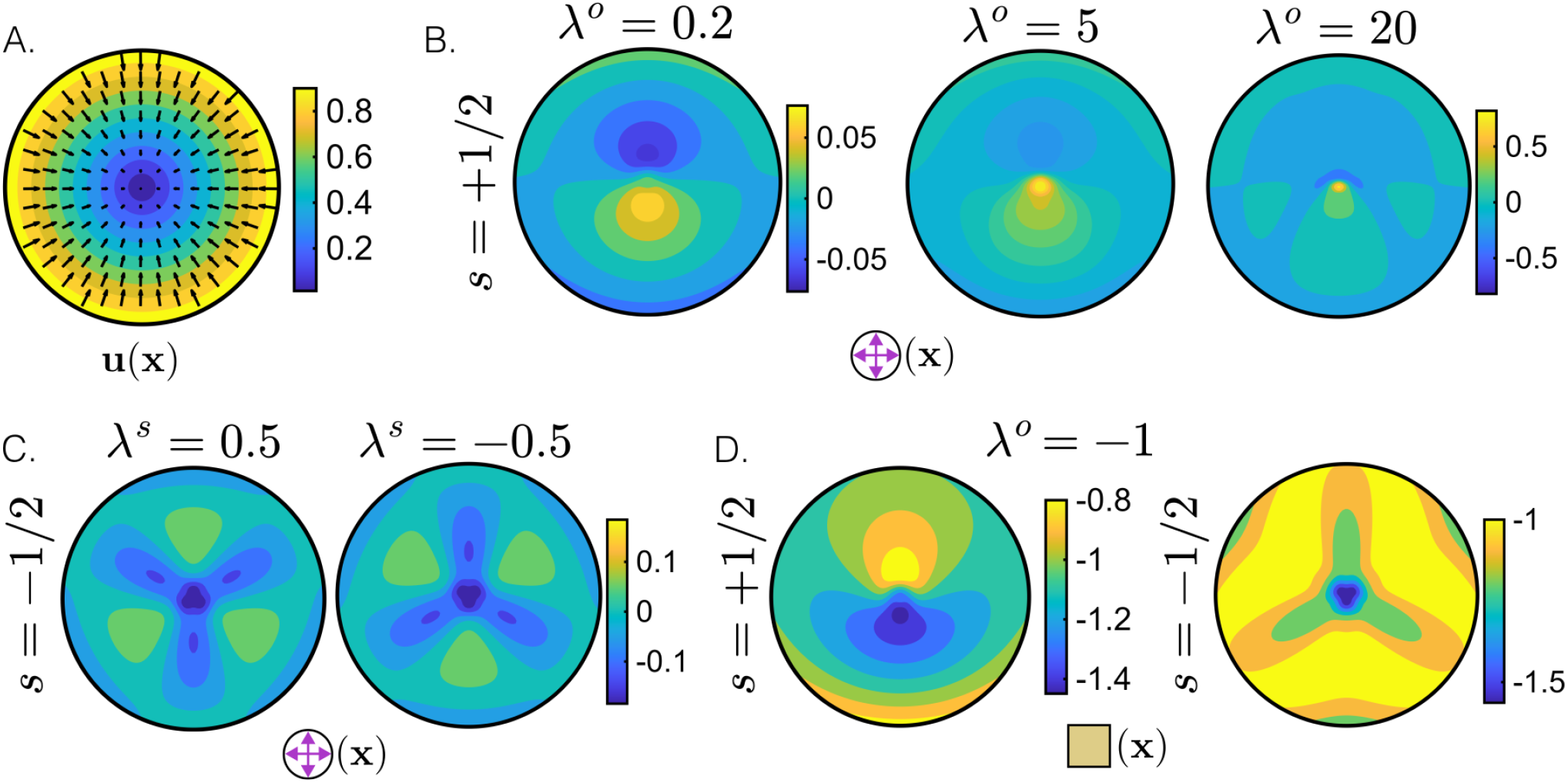
Additional analysis of mechanics around half-integer defects: (A) Steady-state radial flow ***u***(***x***) around defects with charge *s* = +1/2 and −1/2 for parameter sets (i) *δν* = 0.5 or −0.5 and (ii) (*λ*^*s*^, *λ*^*o*^) = (0.5, −0.5) or (−0.5, 0.5). (B) 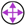(***x***) around a defect for varying *λ*^*o*^ with *λ*^*s*^ = 0. (C) 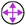(***x***) around a defect for varying *λ*^*s*^ with *λ*^*o*^ = 0. (D) 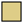(***x***) around defects with (*λ*^*o*^, *λ*^*s*^) = (−1, 0). (A-D) The steady-state mechanical configurations were obtained by solving Eq. (S17). We use the parameter values *m*^0^ = *ν* = 1.

